# Defense by duplication: The relation between phenotypic glyphosate resistance and EPSPS gene copy number variation in *Amaranthus palmeri*

**DOI:** 10.1101/2021.04.13.439524

**Authors:** Sarah B. Yakimowski, Zachary Teitel, Christina M. Caruso

## Abstract

Gene copy number variation (CNV) has been increasingly associated with organismal responses to environmental stress, but we know little about the quantitative relation between CNV and phenotypic variation. In this study we quantify variation in EPSPS (5-enolpyruvylshikimate-3-phosphate synthase) copy number using digital drop PCR and variation in phenotypic glyphosate resistance in 22 populations of *Amaranthus palmeri* (Palmer Amaranth), a range-expanding agricultural weed. Overall, we detected a significant positive relation between population mean copy number and mean resistance. The majority of populations exhibited high glyphosate resistance, yet maintained low-resistance individuals resulting in bimodality in many populations. We investigated linear and threshold models for the relation between copy number and resistance, and found evidence for a threshold of ~15 EPSPS copies: there was a steep increase in resistance before the threshold, followed by a much shallower slope. Moreover, as copy number increases, the range of variation in resistance decreases. This suggests a working hypothesis that as EPSPS copies and dosage increases, negative epistatic interactions may be compensated. We detected a quadratic relation between mean resistance and variation (s.d.) in resistance, consistent with the prediction that as phenotypic resistance increases in populations, stabilizing selection decreases variation in the trait. Finally, patterns of variation across the landscape are consistent with less variation among populations in mean copy number / resistance in Georgia where glyphosate resistance was first detected, and wider variation among populations in resistance and copy number in a more northern state where resistance evolution may be at a younger evolutionary state.

## 1 INTRODUCTION

Variation among individuals in the number of copies of a gene (copy number variation - CNV) arises from gene duplications. Gene copies have long been recognized to play a significant role in genome evolution at the macro-evolutionary scale by providing the opportunity for neo- or sub-functionalization (Force et al., 1999; Lynch & Conery, 2000), or for the eventual loss of a gene copy (Smet et al., 2013). In contrast, the effect of maintaining multiple copies of the same gene on phenotypic variation at the micro-evolutionary scale has received less attention (Hahn, 2009; Zhang, 2003). Increased protein production due to CNV (i.e. gene dosage) can contribute to variation in protein expression (Gaines et al., 2010), conferring a significant effect on the phenotype. If this phenotype increases fitness, gene copies may be maintained (i.e. gene conservation). However, there remains a gap between our knowledge of the molecular mechanisms of gene duplication that result in CNVs, and the evolutionary processes that shape CNV in populations (Hahn, 2009). Therefore, quantifying variation in gene copy number within and among populations, as well as the extent to which gene copy number variation explains variation in key phenotypic traits, are both important steps toward understanding the role of copy number variation in adaptive evolution.

Although many plant housekeeping genes ‘resist’ duplication maintaining a single copy (Smet et al., 2013), genomic studies are increasingly uncovering genes that exhibit CNV (Schrider & Hahn, 2010). The magnitude of detected CNVs varies widely in plants from tens of copies (*Arabidopsis thaliana*; Zmienko et al., 2020) to >100 (*Amaranthus palmeri*; Gaines et al., 2010), to several hundred (*Mimulus guttatus* >600 copies; (Nelson et al., 2019). And, there is increasing evidence that CNV affects phenotypes and fitness, especially for traits involved in responses to environmental stress. For example CNVs have been associated with soil ecotypes in *Arabidopsis lyrata* (Guggisberg et al., 2018), with key life history traits critical for climate adaptation in *Picea glauca* (Prunier et al., 2017) and with decreased flowering time to increase female fitness under drought in *Mimulus guttatus* (Nelson et al., 2019). CNVs have also been associated with stress tolerance and resistance including plant freezing tolerance (Knox et al., 2010), insect resistance to pesticides (Bass & Field, 2011) and disease resistance in plants (Dolatabadian, Patel, Edwards, & Batley, 2017). However, the strength and shape of the quantitative relation between CNV and phenotypic variation remains unknown. Do phenotypes change gradually as copy number increases, or is a threshold copy number associated with a steep shift in phenotype?

The phenotypic effects of CNV are particularly striking in agricultural weedy plant populations that have evolved resistance to herbicides (Gaines, Patterson, & Neve, 2019). Resistance to glyphosate, one of the most commonly used herbicides in North America, is associated with duplication of the (target) gene EPSPS (5-enolpyruvylshikimate-3-phosphate synthase) in at least eight agricultural weedy plant species (Gaines et al., 2019). EPSPS is a gene in the shikimate pathway necessary for the production of aromatic amino acids and other products of the shikimic acid pathway essential to plant function (Herrmann & Weaver, 1999). Glyphosate binds to the enzyme product of the EPSPS gene, blocking EPSPS activity and disrupting plant function (Steinrücken & Amrhein, 1980). Therefore, plants carrying a single EPSPS copy are putatively susceptible to glyphosate. Plants carrying additional copies of EPSPS produce more EPSPS enzyme (Gaines et al., 2010), and individuals that produce sufficient EPSPS protein to bind glyphosate and have remaining functional EPSPS protein for the shikimate pathway to function are ‘resistant’. Therefore, individuals carrying greater numbers of EPSPS copies are predicted to exhibit greater resistance to glyphosate. Indeed, phenotypic resistance is continuous and can be quantified as the proportion of the plant not exhibiting damage following herbicide application (Baucom & Mauricio, 2008a; Dekker & Duke, 1995). Therefore, a key question is whether, for a given glyphosate concentration, there is a gradual linear increase to maximum phenotypic resistance with increasing EPSPS copy number? Or, is there a threshold EPSPS copy number, above which high phenotypic resistance is conferred?

Subsequent to the origin(s) of EPSPS CNV, the evolutionary trajectory of CNV in populations will be shaped by the balance between gene flow and spatially varying selection. The introduction of a high copy number variant to a population not yet containing EPSPS copy number variation could be a critical mode for the spread of resistance across the landscape. Recent gene flow events may be detected as a discontinuous distribution of CNV: very high frequency of single-copy individuals plus relatively low-frequency high-copy number individuals. The relations between EPSPS copy number, resistance, and fitness will determine the strength of selection for increased copy number. Given consistent herbicide stress and genetic variation for resistance, directional selection is predicted to move mean resistance to a level at which all individuals are protected (Baucom & Mauricio, 2008a; Rausher & Simms, 1989). And, genetic variation for herbicide resistance may be reduced by stabilizing selection. Loss of low-copy number variants could erode the potential to prevent more extreme evolution of resistance. However, if herbicide stress is varied in space or time (Norsworthy et al., 2012), there may be opportunity for the maintenance of genetic variation for low herbicide resistance.

Epistatic interactions may complicate the relation between EPSPS CNV and glyphosate resistance. If epistatic genetic variants segregate within populations, this could result in ‘mismatch’: individuals with low EPSPS copy number exhibiting high glyphosate resistance, or individuals with high EPSPS copy number exhibiting low glyphosate resistance. Epistasis also has the potential to cause variation among populations in the relation between copy number and resistance. There is increasing evidence for geographic mosaics of resistance across the landscape (Baucom & Mauricio, 2008b; Kreiner et al., 2019; Kuester, Chang, & Baucom, 2015, Gaines et al. this issue). And, across geographic ranges populations can exhibit genetic differentiation, even in the case of rapid range expansion (Kreiner et al., 2019; Küpper et al., 2018). This suggests that that even if (glyphosate) resistance evolves via the same putative mechanism (ie: CNV) across the geographic range, regions may vary significantly in other genetic loci that exhibit positive or negative epistasis with EPSPS copy number variation. Variation in epistasis could contribute to variation among populations in the relation between EPSPS copy number and glyphosate resistance. For example, positive epistasis could contribute to steeper slope in the relation between copy number and resistance, whereas negative epistatic loci could lower the slope of that relation. Therefore, investigating the extent of ‘mismatch’ between EPSPS copy number and phenotypic resistance within populations, and the consistency, or lack thereof, of this relation among populations is an important step towards understanding the genetic architecture of resistance.

In this study we focus on quantifying variation in phenotypic glyphosate resistance and EPSPS copy number within and among 22 populations of *Amaranthus palmeri* across eastern North America. The geographic distribution of this dioecious plant was restricted to Mexico and the southwest USA prior to the 1990s, and then rapidly spread to agricultural environments throughout the USA, now being present in 29 states (Heap, 2021 [http://weedscience.org]). Resistance to glyphosate was first recorded in 2005 in Georgia (Culpepper, Whitaker, MacRae, & York, 2008) and the association between glyphosate resistance and increased EPSPS gene CNV was reported by Gaines et al. (2010). Given the ongoing geographic range expansion of *A. palmeri* and its threat to crop yield (Ward, Webster, & Steckel, 2013) there is a clear need to understand the rapid evolutionary dynamics of EPSPS CNV and herbicide resistance within and among populations. Here, we specifically investigate the following questions: (1) How variable is phenotypic glyphosate resistance and EPSPS copy number within populations of *Amaranthus palmeri*? Are low-copy number and low-resistance variants maintained within populations that exhibit high mean resistance? (2) Across populations, does mean EPSPS copy number predict phenotypic resistance? (3) Within each population, is there a positive relation between phenotypic resistance and gene copy number? Is the best-fit statistical model for this relation linear? Or, are threshold models which estimate a threshold copy number whereby the slope of the relation differs before and after the threshold a better fit? (4) For both phenotypic resistance and EPSPS copy number is there a significant relation between the population mean and the population variability (s.d.)? Does variability of these traits decrease as the population mean increases?

## 2 METHODS

### 2.1 Samples and glyphosate resistance phenotyping

Seeds from open-pollinated maternal families were sampled in 2016 from 22 populations of *Amaranthus palmeri* occurring in cultivated agricultural fields in Georgia, North Carolina and Illinois USA, spanning ten degrees of latitude and <12 degrees of longitude (Figure 1, Table S1). Seed from 10-34 female plants were collected from each population. In populations with more than 34 females, seeds were sampled by systematically selecting a female for collection every five paces in a straight line from a haphazard point of entry into the field. In populations with 34 or fewer females, seeds were sampled from all females. For each female, entire inflorescences were removed with garden shears and placed in paper bags to be threshed of seeds.

**FIGURE 1.**
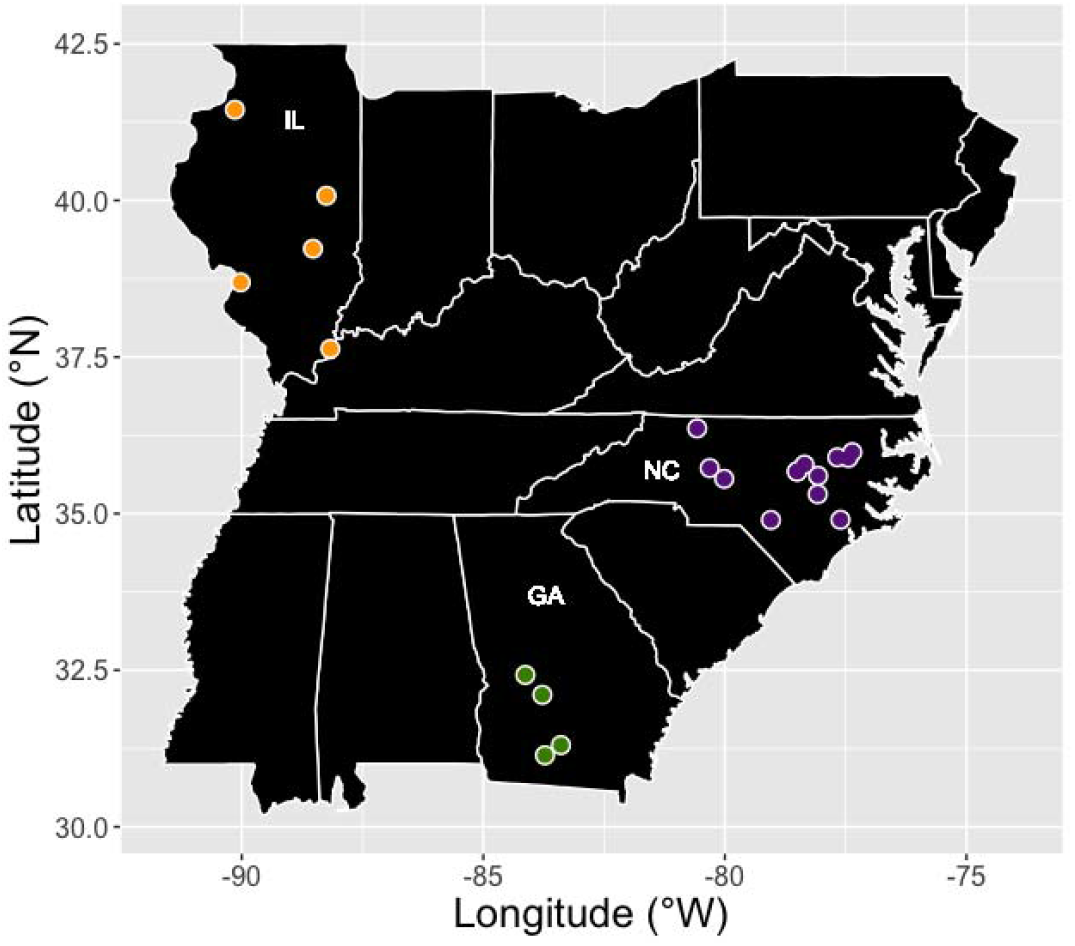
Locations of 22 populations of *Amaranthus palmeri* sampled across eastern North America in Georgia (GA=green), North Carolina (NC=purple), Illinois (IL=orange).

To measure phenotypic resistance, we grew plants from our 22 populations in the University of Guelph Phytotron in 2017. As part of a larger study (Teitel and Caruso, unpublished data), a random sample of families from each population were grown in two temporal blocks (block 1, January - March 2017; block 2, April - May 2017). Families were sampled without replacement, for a mean of 12.18 families per population across both blocks. For each family, we grew 8 plants, resulting in *n* = 2120 plants in block 1 and 2144 plants in block 2. Seeds were germinated in 1.67 L pots. Plants were watered 1-2e daily, fertilized with 50 ml of 17-5-17 fertilizer (Master Plant-Prod Inc., Brampton, Ontario, Canada) every 2 weeks, and exposed to supplemental light (16 h days). The mean temperature and relative humidity (RH) were similar in the two blocks (block 1 = 23.7 °C and 39.0% RH; block 2=23.4 °C and 44.6% RH).

From each family, four randomly-selected plants were glyphosate-treated within a spray chamber in a portable fume hood (Captair^TM^ Flex, Erlab^TM^). Using a hand-held pressurized sprayer attached to 6 mm-diameter tubing with three nozzles, plants were sprayed at the basal rosette stage (mean (1 SD) height = 15.53 (7.12) cm in block 1 and 25.22 (9.61) cm in block 2). Plants were sprayed with a pressure gauge calibrated to 30 psi for 30 s to the point before runoff, such that spray droplets remained on leaves. These glyphosate spraying methods were based on suggested field guidelines, but modified for the contained greenhouse environment. Plants were sprayed with RoundUp^®^ Weathermax (glyphosate, 540 grams acid equivalent per litre, present as potassium salt) at a spray rate of 0.2241 kg a.e. ha^−1^. This spray rate was selected because the next highest rate resulted in ~91% mortality in a preliminary study (Teitel and Caruso, unpublished data).

To estimate phenotypic resistance, we measured leaf damage of glyphosate-treated plants. Glyphosate damage was visually evident as yellowing and shriveling of leaves following glyphosate treatment, and we did not observe any other damage (Teitel, personal observation). Two weeks after glyphosate treatment the number of sprayed leaves with a decaying, yellowing and shriveling pattern were recorded, as well as the total number of leaves on the plant (Baucom & Mauricio, 2008a). This method assumes that new buds were not sprayed. Percent leaf damage (i.e. necrosis) was estimated as the proportion of total leaves damaged following spraying. Phenotypic glyphosate resistance, the proportion of leaves resistant, was calculated as one minus the proportion leaves damaged. The phenotypic resistance data within this manuscript were collected from plants in both blocks, and plants from block 2 were used to estimate EPSPS copy number variation and its relation to resistance (below).

### 2.2 DNA extraction

Prior to glyphosate treatment, one leaf per plant was collected and preserved in silica. DNA was extracted from this silica-dried leaf tissue using the following protocol adapted from Clarke (2009) as follows: We weighed 10 mg of dry tissue and placed it in a 1.5mL micro-centrifuge tube with two UFO-shaped beads. Samples were ground with a tissue homogenizer (Next Advance, Bullet Blender Storm 24) at maximum (12) for 15 seconds to 1 minute, until samples were a fine powder. 500μL of cold CTAB wash buffer was added, mixed, and the mixture was incubated in the fridge for 10 minutes. The mixture was then centrifuged at 10,000g for 2 minutes to pellet tissue. The second phase of wash buffer was then removed and 500 mL of hot CTAB buffer (from a 65°C water bath) was added. Samples were then mixed by inversion and incubated in the 65°C water bath for 30-180 minutes, mixing every 15 minutes. After incubation, we centrifuged tubes at 13,000g for two minutes, removed aqueous layer and placed it into a new tube. To each sample we added 7μL of proteinase K and incubated for 10 mins at 56°C with 300rpm shaking on a hot block. Then we added 5uL RNAse and incubated for 10 minutes at 56°C shaking 300rpm on a hot block. Next we added half-volume guanadine-HCl and half-volume chloroform, mixed immediately and vigorously until thick and cloudy. Samples were centrifuged at 18,000g for two minutes, and the aqueous layer was pipetted into a new 1.5mL tube without disturbing the interphase layer. We added 0.7h volume of 100% isopropanol, mixed samples well by inversion, and incubated samples at room temperature for 10 minutes. Samples were centrifuged at 18,000g for 15 mins, and then isopropanol was carefully poured and/or pipetted off without disturbing the pellet. Pellets were washed in 1mL 70% ethanol, and centrifuged at 18,000g for 2 minutes, 2×. Ethanol (70%) was removed with a pipette and pellets dried in a fume hood for ~10 mins. Pellets were then resuspended in 30μL of nuclease free water, incubated at 56°C for 10 mins to fully dissolve pellet, and then mixed by pipetting.

### 2.3 Digital Drop Estimation of EPSPS copy number variation

#### 2.3.1 Template DNA dilution(s)

The appropriate concentration for DNA samples varies with the copy number of the target gene itself. To multiplex the target gene (EPSPS) with variable CNV and the single-copy reference gene (ALS), the estimates of copies/μL of both target and reference genes need to be above 10 copies/μL to minimize stochasticity, but below ~10,000 copies/μL to ensure sufficient droplets to accurately estimate the ratio. If all droplets are positive, there may be insufficient droplets to accurately estimate the ratio for the sample at that concentration. Therefore, when assaying samples with potentially wide variation in gene copy number, the appropriate DNA template concentration is a balance between being too low for the single copy reference gene (causing high variation among replicates in the ratio of target to reference droplets) and too high for the reference gene (saturating the droplets, and therefore potentially under-estimating the true copy number). Testing ddPCR reactions using DNA template concentrations from 0.01 ng/μL to 1 ng/μL suggested that for the range of variation previously observed in EPSPS in *A. palmeri* (1 to > 140), an initial concentration of 0.7 ng/μl minimized the number of samples with droplet numbers outside the working range described above. When the droplet number for ALS (reference gene) was <10 the reaction was repeated at a higher concentration - 1 or 2ng/μL. When the droplet number for the target gene EPSPS was too high (>10,000, or all droplets saturated = 1,000,000) the reaction was repeated at 0.2 or 0.1 ng/μl, respectively.

#### 2.3.2 Primers and probes for multiplexing amplification of target and reference genes

Amplicons of 75-120bp are recommended to reduce ddPCR error. For amplification of the single-copy reference gene, we used the ALS primer-pair from Gaines et al. (2010) GCTGCTGAAGGCTACGCT and GCGGGACTGAGTCAAGAAGTG in combination with a newly designed probe CGTGCTACTGGACGTGTTGGAGTT 5’ labeled with FAM. This yields a 118 bp amplicon. The primer-pair for amplification of EPSPS was re-designed to yield a smaller amplicon (120bp) than (Gaines et al., 2010; 195 bp amplicon): EPSPS_For: CAAGCTCTCTGGATCGGTTAG and EPSPS_Rev: TCAACATACGGTACGGAAATCA, and was used with EPSPS probe: TGATGGCTACTCCTTTGGGTCTTGG 5’ labeled with HEX. Both probes were double-quenched and contained ZEN/Iowa Black FQ (Integrated DNA Technologies).

#### 2.3.3 Digital drop PCR

The probe-based PCR mastermix included 2xSuperMix (BioRad 186-3023-5) at 1x (10uL per sample), forward target and reference primers at 450 nM each, reverse target and reference primers at 450nM each, target and reference probes at 250nM each, and HindIII (New England Biolabs) (2U, 0.2μL per sample). DNA template (concentration discussed above) and nuclease free water were added to a total volume of 20 μL per reaction. Twenty microlitres of the mastermix + template DNA were pipetted into the very bottom of the central row of (8) wells of the Droplet Generator DG8™ Cartridge (Bio-Rad 186-4008). Sixty-five microlitres of droplet generation oil for probes (BioRad 186-3005) was pipetted into the bottom row of the DG8™ Cartridge, a rubber gasket (BioRad 1863009) was hooked onto the cartridge holder, and then placed in Bio-Rad QX200™ (186-3002) for droplet generation. Upon completion of droplet generation, the gasket was removed and a 50μL multichannel pipette (set at 42μL) was used to very slowly aspirate generated droplet from the top row of the DG8™ Cartridge into the wells of a ddPCR semi-skirted 96-well plate (BioRad 12001925), and droplet generation repeated for all samples as above. Note that a blank negative control water sample was included on each plate. 96-well ddPCR plates were sealed with a pierceable foil (BioRad 1814040) in the plate sealer (Bio-Rad PX1 181-4000) at 180°C for 6 seconds.

C1000 Touch™ thermocycler with deep-well reaction module (BioRad 1851197) was used for the following probe-based amplification: 95°C for 10 mins, followed by 50 cycles of: denaturation at 94°C for 30 sec, annealing at 56°C for 1 min, and elongation at 72°C for 30sec. A final denaturation at 98°C for 10 mins was followed by 10 minutes at 4°C. Finally, the ddPCR plate was placed in the QX200 Droplet Reader (BioRad 1863001) using ddPCR™ Droplet Reader Oil (Bio-Rad 1863004).

Droplet reading was set up in QuantaSoft with the following specifications: CNV1; channel one set to the target (EPSPS HEX-labeled) and channel 2 set to the reference (ALS, FAM-labeled); ABS (absolute readings).

### 2.4 Data Analysis

All data analysis and visualization was performed in R v. 1.2.1335. We tested distributions of phenotypic glyphosate resistance for deviations from unimodality using Hartigan’s drop test (R library “diptest”); a significant deviation from unimodality implies a distribution is at least bimodal. Estimates of skew for distributions of both phenotypic resistance and EPSPS copy number were performed with R library “fBasics”. Variation in phenotypic resistance among populations was investigated with a generalized linear model with a quasibinomial distribution and logit link function. Variation in EPSPS copy number among populations was investigated with a generalized linear model with a poisson distribution. Tukey pairwise contrasts were used to identify which pairs of populations were significantly different for both resistance and copy number.

We investigated the relation between EPSPS copy number and phenotypic glyphosate resistance across the entire data set, and within each of the 22 populations (mean *n* per population=39, *n* range 14-61). We first conducted linear regression. Then, linear two-phase threshold regressions models (Fong, Huang, Gilbert, & Permar, 2017) were used to investigate nonlinear relations between EPSPS copy number and phenotypic glyphosate resistance (R library “chngpt”). Models include a threshold parameter (change point) and examine the relation between copy number and resistance before and after the threshold. We ran ‘upper hinge’ (M10) (Elder & Fong, 2019) and ‘segmented’ (M11) models: ‘upper hinge’ models are most appropriate when there is no significant relation prior to a threshold, whereas the segmented model includes parameters to test for a significant relation before and after the threshold. If more than one of the linear and two threshold models were significant, we compared each pair of significant models using likelihood ratio tests to determine the best-fit model (see Table S4). When analyzing all individuals from all populations together we used a segmented model with population as a random variable. This allowed the model to identify a single threshold EPSPS copy number, but allow populations to vary in their y-intercept of this threshold.

The relation between population mean and population s.d. were examined for both phenotypic resistance and EPSPS copy number using quadratic regression.

## 3 RESULTS

### 3.1 Variation in phenotypic glyphosate resistance

For 22 populations of *Amaranthus palmeri* mean phenotypic glyphosate resistance, estimated as proportion of leaves resistant, ranged from 0.05 to 0.74 (Figure 2) with an overall mean phenotypic resistance of 0.59. Population median resistance ranged from 0 to 0.87. Each population contained some low-resistance (resistance=0-0.1) individuals, but these susceptible individuals varied in frequency from 3 to 40%. The distribution of phenotypic resistance exhibited a deviation from unimodality (Hartigans’ dip test; see Table S2) in 14 of 22 populations. For five of these populations the deviation from unimodality was consistently detected within both of two greenhouse blocks, independently. For eight of these populations the deviation from unimodality was also detected in one of two greenhouse blocks. Five populations exhibit a left-skewed distribution of phenotypic glyphosate resistance (Figure 2, Table S2), and one population (EF-IL) exhibits a right-skewed distribution. Finally, we detected a strong quadratic relation between mean glyphosate resistance and the standard deviation of glyphosate resistance (lm: F_2,19_=34.1, *R*^2^=0.76, P<0.0001) (Figure 6a).

**FIGURE 2.**
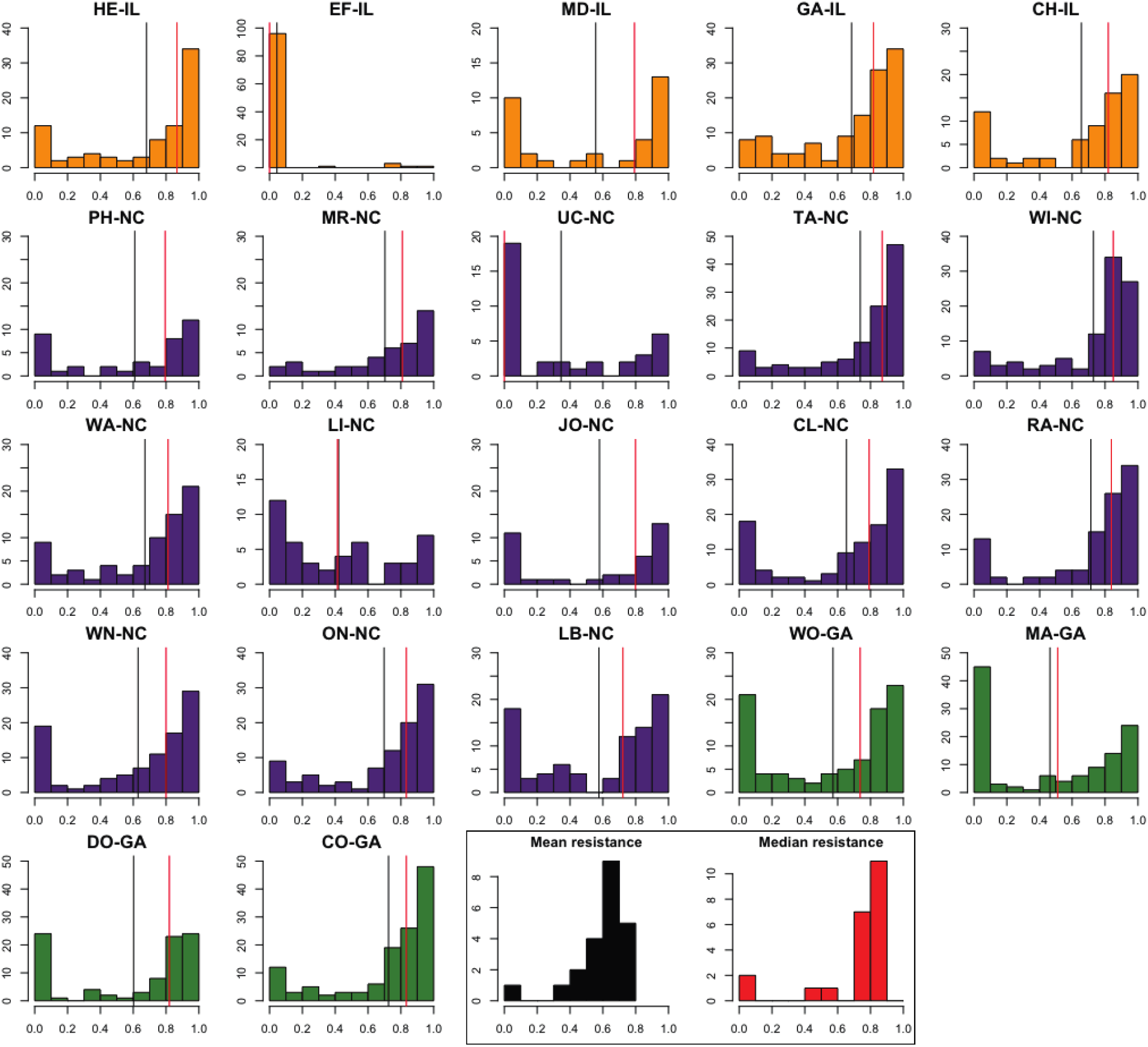
Histograms of the distribution of phenotypic glyphosate resistance in each of 22 populations (ID above) of *Amaranthus palmeri* in eastern North America. Populations are ordered and coloured by state (orange=IL, purple=NC, green=GA). The vertical black and red lines on each histogram represent the mean and median phenotypic glyphosate resistance, respectively. Inset histograms on the bottom row represent the distributions of mean (black) and median (red) glyphosate resistance for our sample of 22 populations.

### 3. 2 Variation in EPSPS gene copy number

Across all 1587 *Amaranthus palmeri* individuals from 22 populations, EPSPS relative copy number ranged from ~1 to 160 (max = 168). Population mean EPSPS copy number ranged from 4.6 to 58.7 and population median EPSPS copy number similarly ranged from 1.03 to ~52.6. Population mean (9 populations) and median (8 populations) copy number most commonly fell within 40-50. The second most-common mean and median EPSPS copy number range was 20-30 copies (7 populations, for both) (Figure 3, inset histograms). Standard deviation (s.d.) of copy number ranges in populations from 14.2 to 29.0. Including the quadratic term increased the fit of the linear model for the regression of the standard deviation of (square-root) EPSPS gene copy number and the mean of (square-root) glyphosate resistance. But although the quadratic term is marginally significant (*t*=−1.9, *P*=0.07), the overall model is not (lm: F_2,19_=2.48, *R*^2^=0.12, *P*=0.11) (Figure 6B).

**FIGURE 3.**
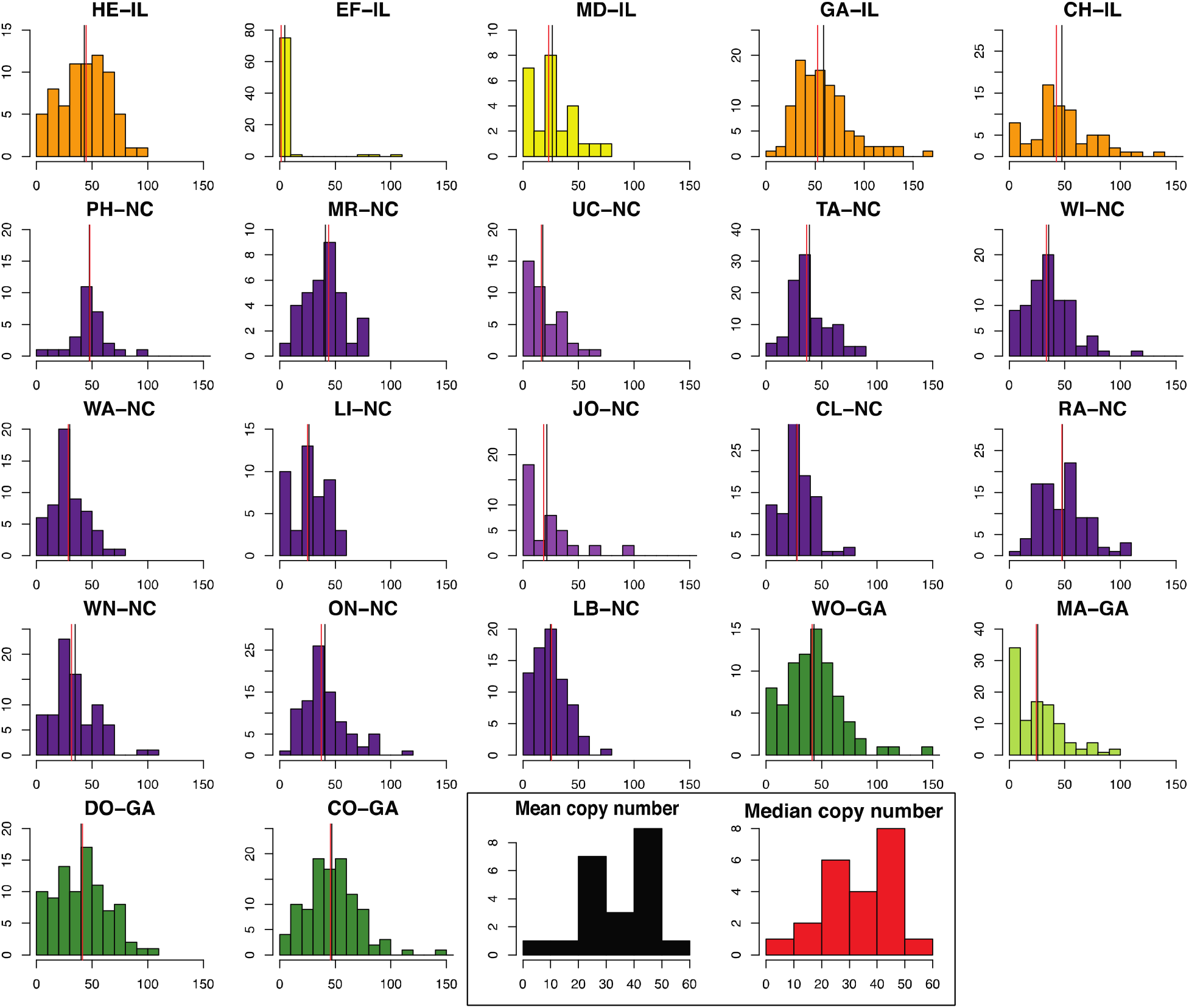
Histograms of the distribution of EPSPS gene copy number variation in each of 22 populations (ID above) of *Amaranthus palmeri* in eastern North America. Populations are ordered and coloured by state (orange=IL, purple=NC, green=GA). Populations for which the observed distribution is strongly skewed right are represented by lighter shades (EF-IL, MD-IL, UC-NC, JO-NC, MA-GA). The vertical black and red lines on each histogram represent the mean and median gene copy numbers, respectively. Inset histograms on the bottom row represent the distributions of mean (black) and median (red) copy number for our sample of 22 populations.

The six most-skewed populations (MA-GA, JO-NC, WI-NC, UC-NC, MD-IL, EF-IL) exhibit distributions of EPSPS copy number strongly skewed to the right with the peak of their distributions at 0 to 10 EPSPS copies. Of these, we only detected individuals carrying >100 gene copies in one population (WI-NC), and at very low frequency. In contrast, the seven populations with the highest kurtosis (>1-30) exhibit long thin tails composed of high copy-number individuals (GA-IL, PH-NC, CO-GA, WO-GA, ON-NC, JO-NC, EF-IL).

Despite the variation in EPSPS copy number within populations, analysis of variation among populations suggests that there is significant variation among populations (glm): specifically, Tukey HSD pairwise suggest significant difference between 103 of 231 pairs of populations (see Table S3). We detected no relation between mean EPSPS copy number and sample size (*R*^2^=0.03, *P*=0.21).

### 3.3 Relation between EPSPS gene copy number and glyphosate resistance

Overall, there is a positive linear relation between glyphosate resistance and EPSPS gene copy number (lm: ß=0.0068, *R*^2^=0.22, *P*<0.0001). However, a segmented threshold model exhibits an even better fit (LRT: *P*<0.0001): there was a significant positive relation with a slope of 0.04 (*P*<0.0001) up to the estimated changepoint in slope at 15.7 EPSPS copies (95% CI: 11.4-20.3) and corresponding to >0.7 resistance (Figure 4B). Following this changepoint, there is a shallower positive relation between EPSPS copy number and resistance with a slope of 0.0012 (*P*<0.0001).

**FIGURE 4.**
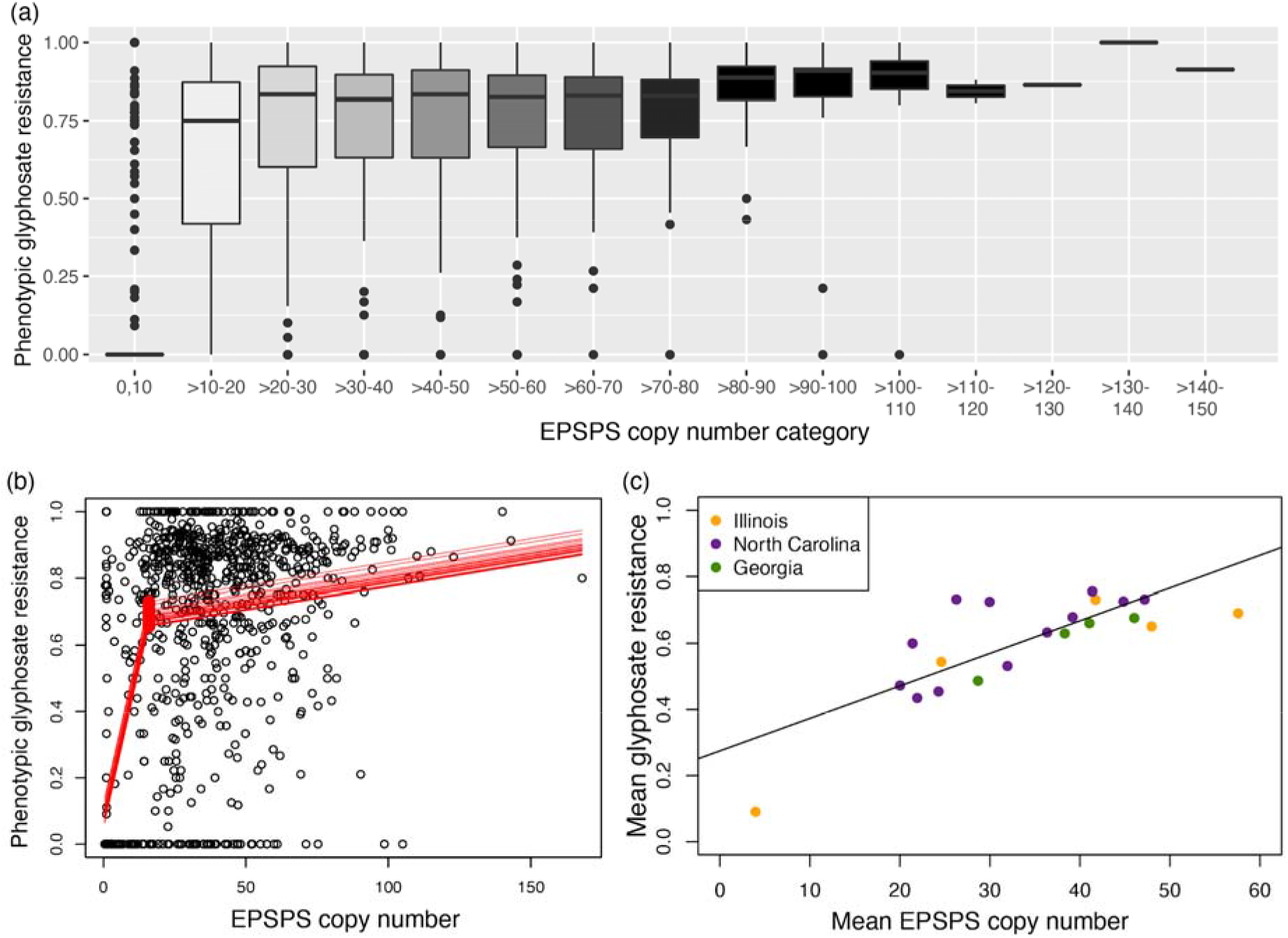
Relation between EPSPS copy number and glyphosate resistance for individuals from 22 populations of *Amaranthus palmeri* in eastern North America. (a) Boxplot of glyphosate resistance for EPSPS copy number visualized by categories of 10 (1-10, >10-20…). The (central) horizontal line of each box represents the median. The lower and upper edge of the boxes represents the 25^th^ and 75^th^ quartiles, respectively. The vertical whiskers extend 1.5x the interquartile range. ‘Outlier’ data points occur beyond the whiskers (> 1.5x the interquartile range). (b) Scatterplot of relation between EPSPS copy number estimated by digital drop PCR and phenotypic glyphosate resistance for >800 plant samples. Fitted lines per population represent the predicted relation from the threshold segmented model (library ‘chngpt’), and estimated EPSPS copy number of ~15 as the threshold value (red points) at which the slope of the relation changes. (c) Scatterplot of mean glyphosate resistance by mean EPSPS gene copy number for 22 populations (orange=IL, purple=NC, green=GA) of *A. palmeri*. The best-fit threshold model fit is shown in red, and linear fit shown with dotted black line.

Visualizing variation in glyphosate resistance by EPSPS gene copy number categories (bins of 10) across all individuals also shows the substantial increase in median resistance from the 0-10 copy category (median resistance = 0) to the >10-20 copy category (median resistance=0.7) (Figure 4A). There was also a reduction in the variation in glyphosate resistance with increasing EPSPS copy number category: the inter-quartile range narrows and fewer outlier individuals are detected. Individuals carrying >10-20 copies exhibited resistance ranging from 0-1, whereas for individuals carrying >80-90 copies resistance ranged from 0.6-1 (box), with only two individual outliers. Finally, although the number of individuals carrying >100 EPSPS copies were relatively few, these individuals consistently exhibit high resistance.

Overall, there was a significant positive linear relation between population mean EPSPS copy number and population mean glyphosate resistance (lm: ß=0.0098, *R*^2^=0.60, *P*<0.0001) (Figure 4C). Note that populations from each the three states (colour=state) are inter-mingled along this relation, with populations from Illinois (orange) exhibiting the broadest range of population mean resistance and mean EPSPS copy number. A segmented threshold model was an even better fit (LRT: *P*=0.002) detected a changepoint at a mean of 26.3 EPSPS copies (95% CI range 21.4-44.8) with a slope of 0.024 (*P*=0.03) prior to the changepoint and no significant relation following the changepoint (ß=0.004, *P*=0.12).

We examined the relation between EPSPS copy number and phenotypic resistance within each population using a linear model and two threshold models (‘upper hinge’, ‘segmented’). A linear model was the best fit model for 5 of the 22 populations (Figure 5, Table S4). The slope of these within-population linear relations ranged from 0.007 to 0.01 (mean ß=0.09); that is an average increase of 9% in resistance for each additional gene copy. The proportion of variation explained by the model (*R*^2^) ranged from 0.13 to 0.46 (Table S4). For 8 of those 22 populations, one or both of the threshold models were a better fit than the linear model (Figure 5, Table S4). For 7 of these populations (all but WN-NC), the linear model was also significant despite not being the best-fit model. For five of the eight populations there was no significant difference between the “upper” and “segmented” models; the “segmented” model only was best fit for two populations, and the “upper” model was best-fit for one population (GA-IL). For the seven significant “segmented” models the estimated threshold EPSPS copy number value ranged from 1.9 to 25.7 copies (mean=13.9).

**FIGURE 5.**
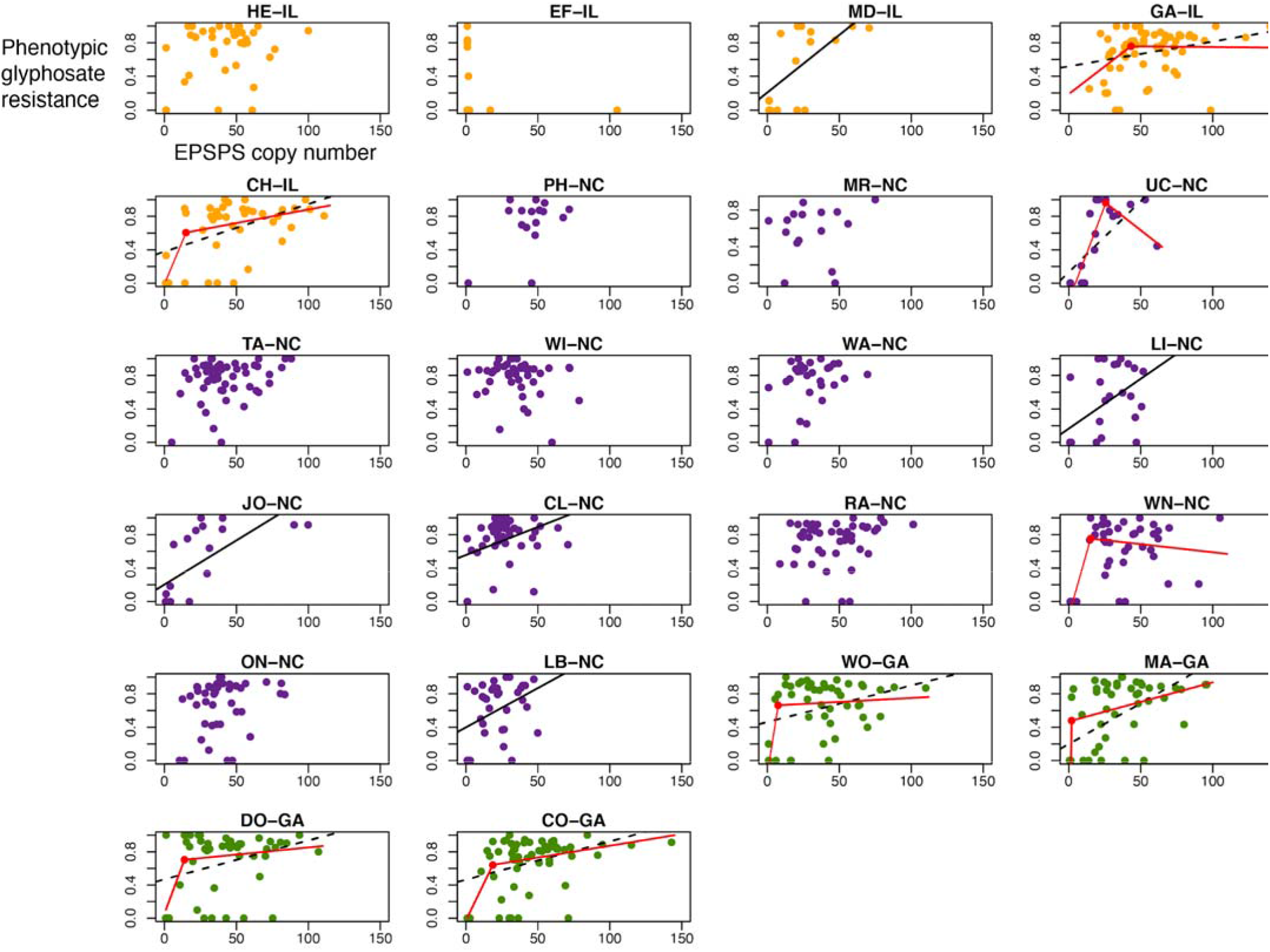
Relation between EPSPS gene copy number and phenotypic glyphosate resistance within each of 22 populations of *Amaranthus palmeri* in eastern North America (orange=IL, purple=NC, green=GA). For populations exhibiting best-fit for the linear model, the fit is a solid black line. For populations exhibiting a best-fit threshold model, the fit is solid red (red line segments and threshold point) and the linear fit with a dotted black line. Populations lacking model lines exhibited no significant relation for all models.

**FIGURE 6.**
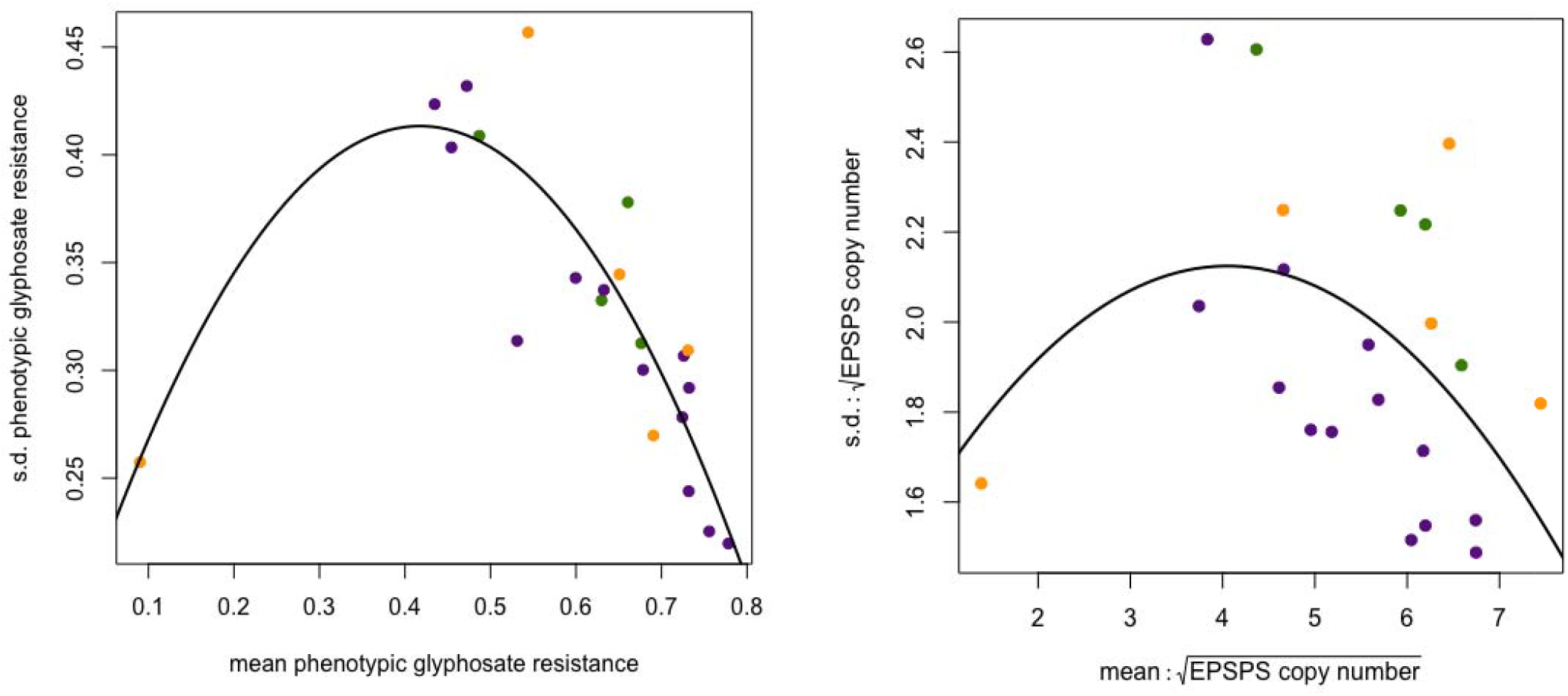
Quadratic relation among 22 populations of *Amaranthus palmeri* (orange=IL, purple=NC, green=GA) for (a) population mean phenotypic glyphosate resistance and populations standard deviation (s.d.) of glyphosate resistance and (b) population mean for square-root copy number and population standard deviation of square-root copy number variation.

Nine populations did not exhibit a significant relation between EPSPS copy number and glyphosate resistance for any of the models examined (Figure 5, Table S4). Note that there was no significant difference in sample size between the populations that best-fit a linear, threshold, or no model (lm: *F*_2_,_19_=2.94 *P*=0.08).

## 4 DISCUSSION

Our investigation of variation in, and the relationship between, phenotypic glyphosate resistance and EPSPS gene copy number variation is consistent with EPSPS copy number variation playing a significant role in glyphosate resistance across eastern populations of *Amaranthus palmeri*. Within populations, phenotypic glyphosate resistance exhibited a bimodal distribution in 14 populations. And, low-copy number variation in EPSPS was present within populations, even in populations with individuals carrying >100 EPSPS copies. Among populations mean EPSPS copy number variation explains ~57% of variation in mean phenotypic glyphosate resistance. Overall, we detected a threshold EPSPS copy number of ~15, above which phenotypic glyphosate resistance is on average above 0.7. This is followed by a gradual increase in phenotypic resistance and reduction in the frequency of low-resistance phenotypes with increasing EPSPS gene copy number. Finally, there was a significant quadratic relation between mean phenotypic resistance and s.d. phenotypic resistance, consistent with the prediction that stabilizing selection may decrease variation in resistance as mean resistance increases. Here we discuss the demographic and evolutionary factors that may contribute to observed patterns of variation in both phenotypic glyphosate resistance and EPSPS CNV, and how this variation may inform future study of the evolutionary trajectory of resistance in this range-expanding agricultural weed.

### 4.1 Maintenance of variation in phenotypic glyphosate resistance

Overall mean phenotypic glyphosate resistance was high, yet all populations of *Amaranthus palmeri* retained some individuals exhibiting low-resistance. Moreover, variation in phenotypic glyphosate resistance exhibited bimodal distributions, with peaks at the two extremes of phenotypic resistance, in up to 14 of the 22 populations studied. To our knowledge, bimodal distributions of herbicide resistance within populations have not been previously reported. The occurrence of low-resistance individuals are significant to pest management: if low-resistance individuals have the opportunity to mate with higher-resistance individuals (Winterer & Weis, 2004), this may contribute to the maintenance of phenotypic variation in glyphosate resistance, preventing high-resistance from being fixed in a population. Moreover, it raises the question of what environmental or genetic factor(s) have contributed to the maintenance of low-resistance individuals in high-resistance populations.

The balance of gene flow among populations vs. the magnitude of glyphosate selection within populations may contribute to the bimodality of phenotypic glyphosate resistance: gene flow from low-glyphosate-resistance sources that is frequent enough relative to the strength of glyphosate selection could contribute to the maintenance of low-resistance phenotypes. In this study the role of gene flow in the introduction of copy number variation to a low-resistance population is observed (Figure 3, EF-IL). However, to estimate the role of ongoing gene flow in the maintenance of low-resistance variation future study of population genetic structure (Kreiner et al., 2019; Küpper et al., 2018) combined with glyphosate resistance phenotyping at a fine spatial scale and with contiguous sampling of populations is needed. This type of study would also contribute to estimating the frequency of agricultural environments that harbour low-resistance individuals and the role of gene flow in maintaining variation in glyphosate resistance. Overall the use of glyphosate has been sustained or even continued to increase in the United States over the past decade (Benbrook, 2016; Duke, 2018), with growers of Glyphosate-Resistant crops are increasingly relying on glyphosate rather than other herbicides (Duke, 2018; Givens et al., 2009). Therefore, these environments may select for increasingly lower frequency of low-resistance. Agricultural environments that produce non-GR crops and/or organic agricultural environments may harbour higher frequencies of low-resistance individuals. Moreover, future estimates of glyphosate resistance in agricultural environments lacking glyphosate selection could provide insight into the rate of gene flow across the landscape, and also whether EPSPS copy number variation persists in populations that are selectively neutral, with respect to glyphosate stress.

One environmental factor that could contribute to the maintenance of low-resistance individuals is variation in glyphosate stress across the growing season or among years. Germination of *A. palmeri* occurs in multiple cohorts throughout the growing season (Neve, Norsworthy, Smith, & Zelaya, 2011) and plants can also reproduce quickly and at small plant size under crowded conditions (Yakimowski, *personal observation*). This reproductive ability could increase the survival and reproduction of low-resistance individuals between glyphosate applications, contributing to the maintenance of low-resistance individuals in populations of *A. palmeri*. Further, is it possible for populations to maintain cohorts adapted to both non-glyphosate and glyphosate-stressed environments? This could be akin to bimodal distributions arising due to ecological adaptive evolution with divergent selection between conspecific groups that occupy different environments (Anderson, Pauw, Cole, & Barrett, 2016; Hendry, Huber, De León, Herrel, & Podos, 2009). However, it is important to note that although herbicide treatment would be expected to promote strong assortative mating (Cordeiro, Correa, Rosi-Denadai, Tome, & Guedes, 2017; Doebeli, 1996) between high-resistance individuals, in the absence of glyphosate stress intermating between low- and high-resistance individuals is expected. Therefore, low-resistance individuals would be predicted to decrease in frequency over time, unless differences in phenology (Baucom, 2019; Bonner, Sokolov, Westover, Ho, & Weis, 2019; Winterer & Weis, 2004) between low- and high-resistance phenotypes existed. Indeed, Teitel and Caruso (*unpublished data*) observed variation that earlier emergence was correlated with higher glyphosate resistance. Association between high-resistance and reduced fitness could also contribute to the maintenance of low-resistance individuals at a higher frequency. Investigation of the relation between EPSPS copy number (up to ~65 copies) and fitness found no relation (Vila◻JAiub, Neve, & Powles, 2009). However, Teitel and Caruso (*unpublished data*) observed selection against the individuals exhibiting the highest phenotypic resistance in a high-competition crop environment, but not in a low-competition no-crop environment. Below we also consider another non-mutually exclusive hypothesis that negative epistatic interactions between other loci with EPSPS CNV could contribute to the maintenance of low-resistance phenotypes.

### 4.2 Relation between phenotypic glyphosate resistance and EPSPS copy number variation: towards a framework for understanding evolutionary trajectory

The overall support for a threshold relation between EPSPS copy number and glyphosate resistance suggests that the relation between EPSPS copy number and glyphosate resistance varies over the observed range of copy number variation. The full threshold model (Figure 4B) suggests that an accumulation of ~15 EPSPS copies predicts a steep increase in phenotypic resistance to ~0.65-0.75. Subsequent acquisition of >15 copies is associated with a shallow increase in resistance of ~0.12% per copy. The better fit of the threshold model relative to the linear regression model suggests that the selection gradient around this threshold could be very strong, if resistance is strongly correlated with fitness (Moorad & Wade, 2013). Therefore, under glyphosate stress, and with EPSPS copy number variation available, copy number variants 2-15 would be predicted to rapidly increase in frequency. However, beyond this threshold, subsequent evolution of resistance may proceed at a much slower rate. Yet, if the generation of variation in EPSPS copy number increases, this variability may counter the shallower selection gradient. Unequal segregation of the extrachromosomal DNA EPSPS copies (Koo et al., 2018) may rapidly increase heterogeneity of copy number variation (Turner et al., 2017). Overall this suggests that the presence of even low EPSPS CNV in combination with glyphosate selection predicts rapid evolution of phenotypic resistance.

What is driving the evolution of EPSPS copy number variation to population means of ~58 and individual copy numbers upwards of 160, if substantial resistance is observed with ~15 copies? One potential factor could be negative epistatic interactions between EPSPS CNV and other genetic loci contributing to glyphosate resistance. Variation among individuals in phenotypic glyphosate resistance declines with EPSPS copy number (Figure 4A): at moderate EPSPS copy numbers (20-70) median resistance is high, but there are individuals that exhibit low to moderate resistance. This pattern suggests there may be variants segregating within populations that negatively interact with EPSPS CNV, countering the increase in resistance conferred by EPSPS copy number and lowering phenotypic resistance in these individuals. As EPSPS copy number increases upwards of 100, these ‘mismatched’ high-copy number, low resistance individuals become rarer. One possibility is that as EPSPS copy number increases the increased dosage effect gradually overwhelms negative epistatic effects. Therefore, this predicts that as mean copy number increases, the variability in copy number and resistance should decrease.

Stabilizing selection is predicted to decrease variation around the trait mean. Although only one of our populations exhibited very low mean resistance and copy number, we detected a quadratic relation between mean resistance and s.d. resistance (Fig 6A). This relation is consistent with the early stages of glyphosate resistance evolution being associated with increasing variation in resistance, followed by a decline in variation, consistent with stabilizing selection, as mean resistance increases. We observe a similar, albeit not statistically significant pattern for EPSPS copy number mean and s.d. (Fig 6B). Stabilizing selection would also predict that fitness of individuals at phenotypic extremes would be lower, and should be investigated in future work. If extreme high copy number individuals exhibit lower fitness (Martin et al., 2017) this will suggest that the current resistance variation in the population may stabilize. However, if lower fitness is not lower in high copy number individuals, there may remain further potential for evolution of increased resistance.

Within each of the geographic areas (states) studied we observe a range of mean resistance and mean EPSPS copy number, rather than a clustering by geographic region (Figures 4C & 6). The most northern region included in this study, Illinois, may have been the most recent to evolve glyphosate resistance (Davis, Schutte, Hager, & Young, 2015). Notably, populations in this region exhibit the widest variation in mean resistance and mean copy number among the (five) populations studied here, beyond the range of variation observed among populations in either Georgia or North Carolina (Figure 4C). This suggests that as *A. palmeri* expands its range northward, variation in resistance among populations may be high. Variation among populations that have been more recently established and are early in their evolutionary trajectory of resistance could be the result of genetic bottlenecks associated with founder effects (Baker & Moeed, 1987). But, as resistance and EPSPS CNV spreads across the landscape and a balance between gene flow and selection for resistance is established, variation both within and among populations may decline. Indeed, our data are consistent with this hypothesis showing the least variation in mean resistance / copy number among the Georgia populations, the state in which resistance to glyphosate was first recorded. Greater sampling of populations combined with historical records of introduction and emergence of glyphosate resistance are needed to test this landscape level hypothesis.

### 4.3 Variation in the EPSPS copy number - resistance relation

In *Amaranthus tuberculatus* there is some evidence of alternative mechanism(s) for glyphosate resistance: two populations exhibiting glyphosate resistance were found to exhibit no or very little EPSPS CNV (Chatham et al., 2015), and altered translocation/uptake was a suggested alternative mechanism (Nandula, Ray, Ribeiro, Pan, & Reddy, 2013). In contrast, overall we found little evidence of alternative glyphosate mechanisms in this study. Of the 105 individuals exhibiting EPSPS copy number between 0.4 and 1, all but seven individuals exhibited zero resistance. And, of 45 individuals with copy number between 1 and 2, seven individuals exhibited resistance ranging 0.1 to 1. However, it is possible there are loci that exhibit positive epistasis with EPSPS CNV, and contribute to the high resistance phenotypes just beyond the threshold of 15. For example, in WO-GA and MA-GA both have slope of segment 1 much higher and an estimated threshold of 2 copies. These populations may be future candidates for investigation of traits that exhibit positive epistasis with EPSPS.

Negative epistatic interactions with EPSPS CNV have the potential to decrease the phenotypic effect of EPSPS copy number. In studies of microbial resistance negative epistasis has been frequently detected (Brown et al., 2020; Lukacisinova, Fernando, & Bollenbach, 2020; Yang et al., 2020). One way we were able to look at this was by examining the ratio of the frequency of low EPSPS copy number (<=10) relative to the frequency of low phenotypic resistance (0-0.1). For most populations with a ratio close to 1, low copy number variants likely explain the frequency of low-resistance individuals. However, many populations exhibited a ratio much lower than one (Figure S1), suggesting that the frequency of low copy number individuals alone may not account for the frequency of low-resistance individuals. GA-IL was one of the four populations with the largest deviation from 1, and this was also the only population that exhibited a better fit to the upper threshold model (than segmented), with no significant relation between resistance and EPSPS copy number prior to the much-higher threshold of 43 copies estimated within this population. Populations exhibiting the greatest frequency of high-copy number to low resistance ‘mismatch’ and/or deviation from the ratio of one are good candidate populations for studying negative interactions between EPSPS copy number and other genetic loci. Epistatic interactions among microbial resistance mechanisms is common (Hall & MacLean, 2011; Sousa, Balbontín, Durão, & Gordo, 2017). Given reports of resistance to at least four other chemical classes herbicide, and multiple resistance, in *A. palmeri* (Ward et al., 2013), future investigation of interaction among the genetic loci underlying these resistance mechanisms seems warranted.

## 5 Conclusion

We investigated variation in, and the relation between, phenotypic glyphosate resistance and EPSPS copy number in the range-expanding weed *Amaranthus palmeri*. We detected a significant threshold in herbicide resistance at ~15 EPSPS copies. Yet, we also detected a surprising amount of low-resistance phenotypic variation standing in populations. On one hand this suggests potential for management strategies that prevent the fixation of high-resistance. On the other hand, variation in the strength and direction of glyphosate selection, a management strategy meant to slow the evolution of resistance, may also have the potential to increase variation and create more starting points for evolution to navigate adaptive peaks (Steinberg & Ostermeier, 2016). Therefore, it will be critical for future work to focus on understanding the fitness effects of high resistance and high EPSPS copy number. Moreover, investigation of the evolutionary dynamics between glyphosate resistance, other stress resistance mechanisms, and life history traits for populations across the expanding range of *A. palmeri* is called for.

## ACKNOWLEDGEMENTS

We thank David Jordan, Andrew Hare, Stanley Culpepper, Wes Everman, Patrick Tranel, Jeremy Kichler, Ronnie Barentine, Kurt Maertens, Eric Alinger, Kevin Johnson, Brian Schoon, Nikki Keitner, Lanae Ringler, and Larry Uthell for help sampling populations, Michael Mucci for building glyphosate spray chamber, and Tannis Slimmon, Ann Lee, Sarah McDonald, Aaron Hudson, Emily Williams, Jennifer Wood, Jasmin Dawson, Lucy Burns, Chloe Katsademas for assistance with resistance phenotyping. We appreciate the work of Lisa Han, Graeme McLeod and Zhengxin Sun with the ddPCR assays. We thank the Tayade Lab (Department of Biomedical Sciences, Queen’s U) for their generous access to BioRad digital droplet platform. This work was funded by an NSERC Discovery grant to C.M.C., an NSERC Alexander Graham Bell CGS - D to Z.T., and an NSERC Banting Fellowship to S.B.Y.

## DATA ACCESSIBILITY

Data will be made available on Dryad.

## AUTHOR CONTRIBUTIONS

S.B.Y, Z.T and C.M.C. designed this study; Z.T. conducted the glyphosate resistance phenotyping. S.B.Y. conducted ddPCR assays of EPSPS copy number variation. S.B.Y analyzed data and wrote paper with comments and edits from Z.T. and C.M.C.

