## Supplements for "Defense by duplication: The relation between phenotypic glyphosate resistance and EPSPS gene copy number variation in *Amaranthus palmeri*"

Supporting Information.

TABLE S1. Locations of the 22 populations of *Amaranthus palmeri* in eastern North America. The last two letters of the ‘Population ID’ correspond to state (IL=Illinois, NC=North Carolina, GA=Georgia).

| **Population** | **Locality** | **Latitude (°N)** | **Longitude (°W)** |
| --- | --- | --- | --- |
| HE-IL | Henry Co. | 41.447092 | -90.141592 |
| CH-IL | Champaign Co. | 40.074505 | -88.246533 |
| EF-IL | Effingham Co. | 39.228864 | -88.525873 |
| MD-IL | Madison Co. | 38.691638 | -90.021769 |
| GA-IL | Gallatin Co. | 37.629676 | -88.166345 |
| PH-NC | Pine Hill | 36.360356 | -80.57341 |
| MR-NC | Martin Co. | 35.979622 | -77.362871 |
| UC-NC | Upper Coastal Plain | 35.897903 | -77.672789 |
|  | Research Station |  |  |
| TA-NC | Tarboro | 35.887358 | -77.443634 |
| WA-NC | Wake Co. | 35.781702 | -78.352149 |
| LI-NC | Linwood | 35.726837 | -80.313716 |
| CL-NC | Clayton | 35.674307 | -78.512462 |
| JO-NC | Johnston Co. | 35.669931 | -78.49939 |
| WI-NC | Wilson Co. | 35.60278 | -78.071049 |
| RA-NC | Randolph Co. | 35.55501 | -80.013807 |
| WN-NC | Wayne Co. | 35.310539 | -78.078181 |
| ON-NC | Onslow Co. | 34.90157 | -77.607704 |
| LB-NC | Lumber Bridge | 34.897865 | -79.047614 |
| MA-GA | Macon Co. | 32.423982 | -84.129029 |
| DO-GA | Dooly Co. | 32.106095 | -83.777492 |
| WO-GA | Worth Co. | 31.3032 | -83.3932 |
| CO-GA | Colquitt Co. | 31.141031 | -83.723125 |

TABLE S2. Summary statistics for variation in phenotypic glyphosate resistance and EPSPS copy number number for 22 populations of *Amaranthus palmeri*.

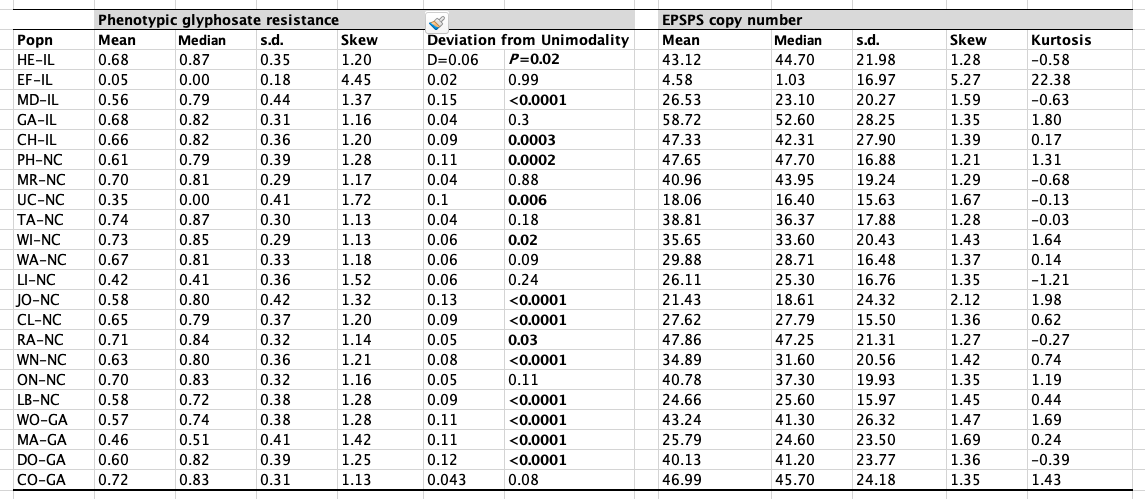

TABLE S3. TukeyHSD pairwise contrasts between 22 populations of *Amaranthus palmeri* for (a) phenotypic glyphosate resistance and (b) EPSPS copy number.

(a)

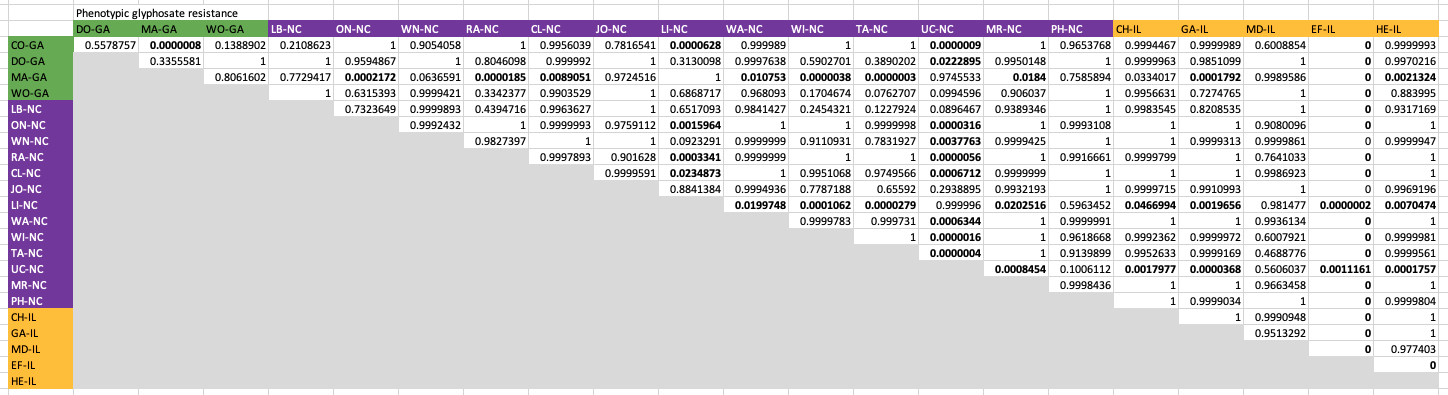

(b)

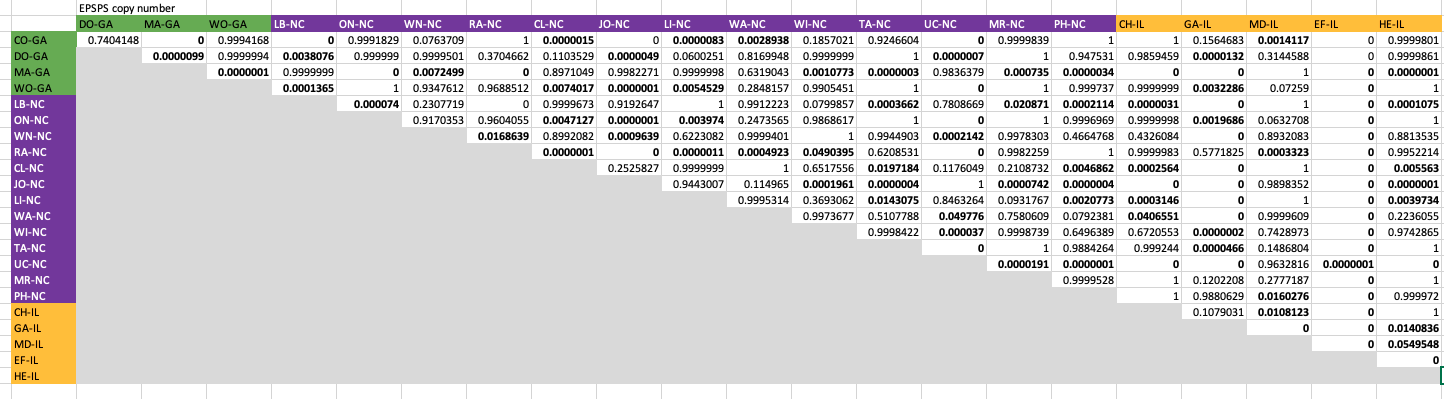

TABLE S4. Linear, ‘upper’ threshold, and ‘segmented’ threshold models for the relation between EPSPS copy number and phenotypic glyphosate resistance for 22 populations of *Amaranthus palmeri.* Significant model fits correspond to the models shown in Figure 5.

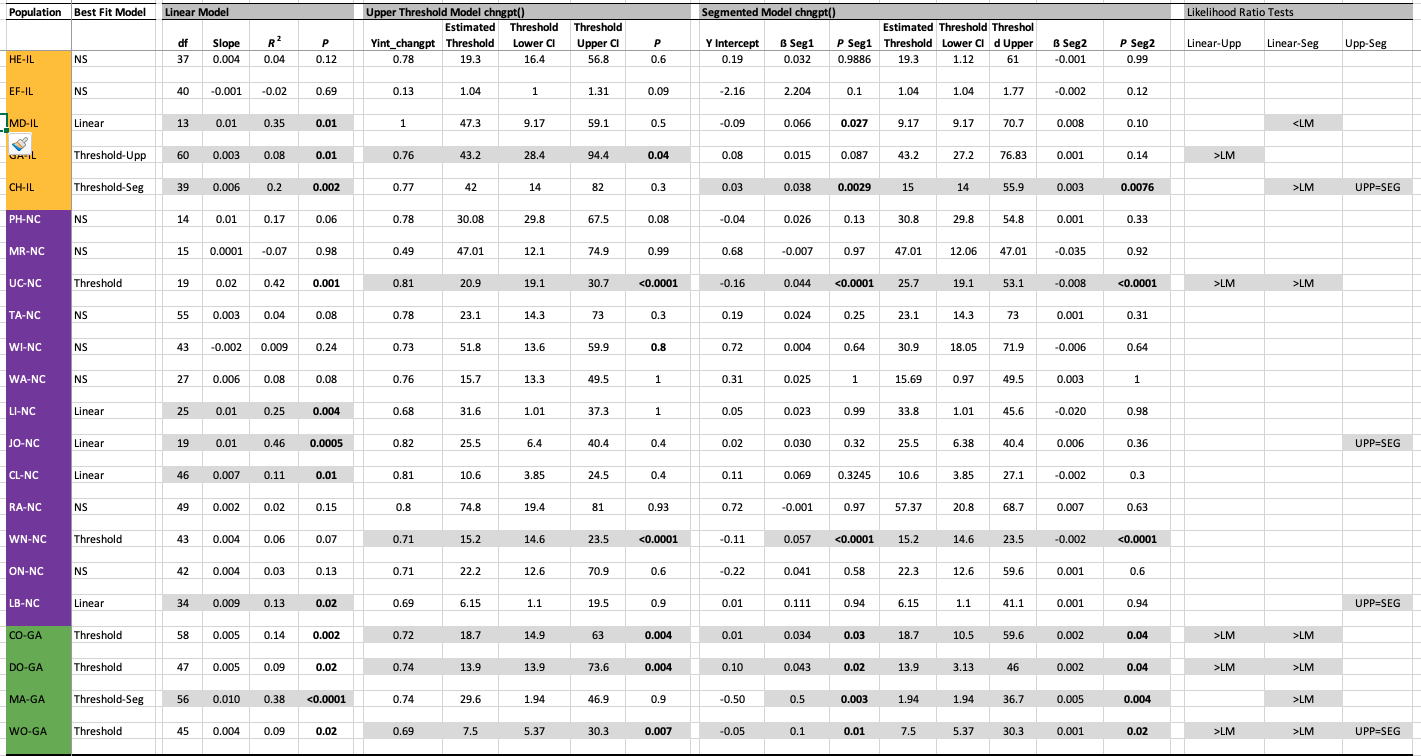

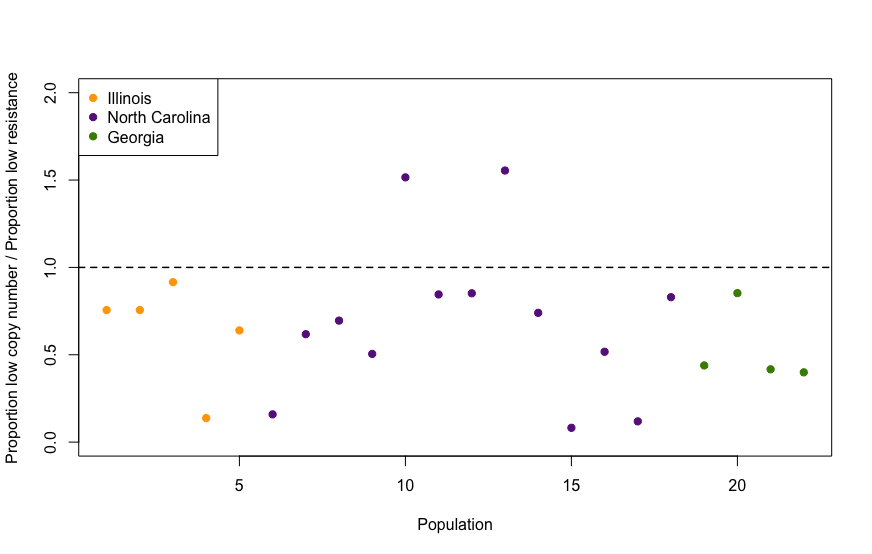

FIGURE S1. The ratio of the frequency of low EPSPS copy number (1-10 copies) individuals to the frequency of low glyphosate resistance (0-0.1) individuals for 22 populations of *Amaranthus palmeri* in Illinois (orange), North Carolina (purple), and Georgia (green). The dotted horizontal line at 1.0 an equal ratio such that low EPSPS copy number accounts for the low-resistance variants in a population. Populations that exhibit values <1 and approaching 0 represent populations for which the frequency of low EPSPS copy number does not account for the frequency of low phenotypic glyphosate resistance.
